# Effects of sub-threshold transcutaneous auricular vagus nerve stimulation on cerebral blood flow

**DOI:** 10.1101/2021.07.13.451709

**Authors:** Conan Chen, Yixiang Mao, Maryam Falahpour, Kelly H. MacNiven, Gary Heit, Vivek Sharma, Konstantinos Alataris, Thomas T. Liu

**Author notes:** **Corresponding Authors:** Conan Chen and Thomas T. Liu. **Present/Permanent Address:** 9500 Gilman Drive #0677, La Jolla, CA 92093.

## Abstract

**Background:** Transcutaneous auricular vagus nerve stimulation (taVNS) has shown promise as a non-invasive alternative to vagus nerve stimulation (VNS) with implantable devices, which has been used to treat drug-resistant epilepsy and treatment-resistant depression. Prior work has used functional MRI to investigate the brain response to taVNS, and more recent work has also demonstrated potential therapeutic effects of high-frequency sub-threshold taVNS in rheumatoid arthritis. However, no studies to date have measured the effects of high-frequency sub-threshold taVNS on cerebral blood flow (CBF).

**Objective/Hypothesis:** The objective of this study was to determine whether high-frequency (20 kHz) sub-threshold taVNS induces significant changes in CBF.

**Methods:** Arterial spin labeling (ASL) MRI scans were performed on 20 healthy subjects in a single-blind placebo-controlled repeated measures experimental design. The ASL scans were performed before and after 15 minutes of either sub-threshold taVNS treatment or a sham control.

**Results:** taVNS induced significant changes in CBF in the superior posterior cerebellum that were largely localized to bilateral Crus I and Crus II. Post hoc analyses showed that the changes were driven by a treatment-related decrease in CBF.

**Conclusions:** Fifteen minutes of high-frequency sub-threshold taVNS can induce sustained CBF decreases in the bilateral posterior cerebellum in a cohort of healthy subjects. This study lays the foundation for future studies in clinical popluations to assess whether similar effects can be observed and are related to treatment outcomes.

## Introduction

Vagus nerve stimulation (VNS) with implantable devices is currently used to treat drug-resistant epilepsy and treatment-resistant depression. However the cost and complications associated with implantation have limited the application of invasive VNS to a broader range of health conditions and spurred growing interest in non-invasive approaches, such as transcutaneous auricular vagus nerve stimulation (taVNS) [1–4]. In taVNS, an electrode on the surface of the ear delivers a small electrical current across the skin to the auricular branch of the vagus nerve. Stimulation of the auricular branch in turn activates other areas of the brain through connected pathways [5,6], including regions involved in the regulation of the autonomic nervous system (ANS). The theory behind the therapeutic efficacy of taVNS is that stimulating the vagus nerve can correct maladaptive processing within the nervous system as well as neuro-immune communications that are associated with negative health outcomes. Building upon prior work that suggests a link between ANS dysfunction and inflammation [7], a recent open-label study of subjects with rheumatoid arthritis demonstrated clinically relevant changes in both disease biomarkers and symptoms after 12 weeks of sub-threshold taVNS treatment [8]. The therapeutic benefits of taVNS have also been reported in the context of depression [1,7,9] and migraine [10], with mounting evidence linking these clinical effects to neural responses [2,3,11].

While the mechanisms through which taVNS causes changes are not completely understood, a number of recent studies have used blood oxygen-level dependent (BOLD) functional MRI (fMRI) to characterize the acute brain response to taVNS [2,3,11–14]. While BOLD fMRI is an effective approach for measuring evoked brain responses, its susceptibility to low-frequency noise sources makes it less effective for measuring changes that occur with time periods greater than several minutes [15]. In contrast, measures of cerebral blood flow (CBF), a well-defined physiological quantity, can be used to characterize changes in brain physiology occurring over periods ranging from seconds to years [16], and therefore constitute a promising metric for the assessment of the sustained effects of VNS.

In prior work, both SPECT and PET have been used to characterize the CBF response to VNS applied using implantable devices in patients with either major depression (MD) or epilepsy. In MD patients, SPECT has been used to demonstrate regionally dependent increases and decreases in CBF due to 4 to 10 weeks of chronic VNS [17,18]. In a PET study with MD patients, Conway et al [19,20] reported regionally dependent increases and decreases in the evoked CBF response to short duration (90s) VNS. For epilepsy patients, Henry et al [21,22] reported PET-based CBF increases and decreases in the evoked CBF reponse to 30s of VNS, with the effects diminishing after 12 weeks of chronic VNS. In addition, two earlier pilot studies [23,24] reported VNS evoked changes in PET CBF measures.

To the best of our knowledge, no studies to date have examined the CBF response to taVNS. In this study, we used arterial spin labeling (ASL) MRI to measure changes in CBF induced by 15 minutes of taVNS in healthy subjects, employing a stimulus similar to that of [8]. In contrast to SPECT and PET, ASL MRI does not require the injection of radioactive tracers, making it especially suitable for repeated measures in a healthy population [25,26]. Furthermore, in this study the taVNS is applied at a sub-threshold level, such that subjects cannot readily determine if they are receiving the treatment or a placebo. In contrast to prior studies where the perceptible sensations associated with VNS may have complicated the interpretation of the results, the sub-threshold stimulus used in this study removes somatosensory confounds and is therefore compatible with a single-blind placebo-controlled design. The goal of this study was to determine whether a single course of taVNS with duration similar to that of the daily dose employed in [8] could lead to detectable changes in CBF.

## Material and methods

### Overview

We used a single-blind placebo-controlled experimental design. Twenty seven healthy subjects were enrolled in this study after providing informed consent (14 females and 13 males, aged 19-40 years). The study was approved by the UCSD Institutional Review Board (IRB), and informed consent was obtained from all participants. Figure 1 shows the overall experiment structure. Each subject participated in a control session and a treatment session, with the order randomized across subjects. The sessions were separated by at least one day to avoid taVNS carryover effects.

**Figure 1:**
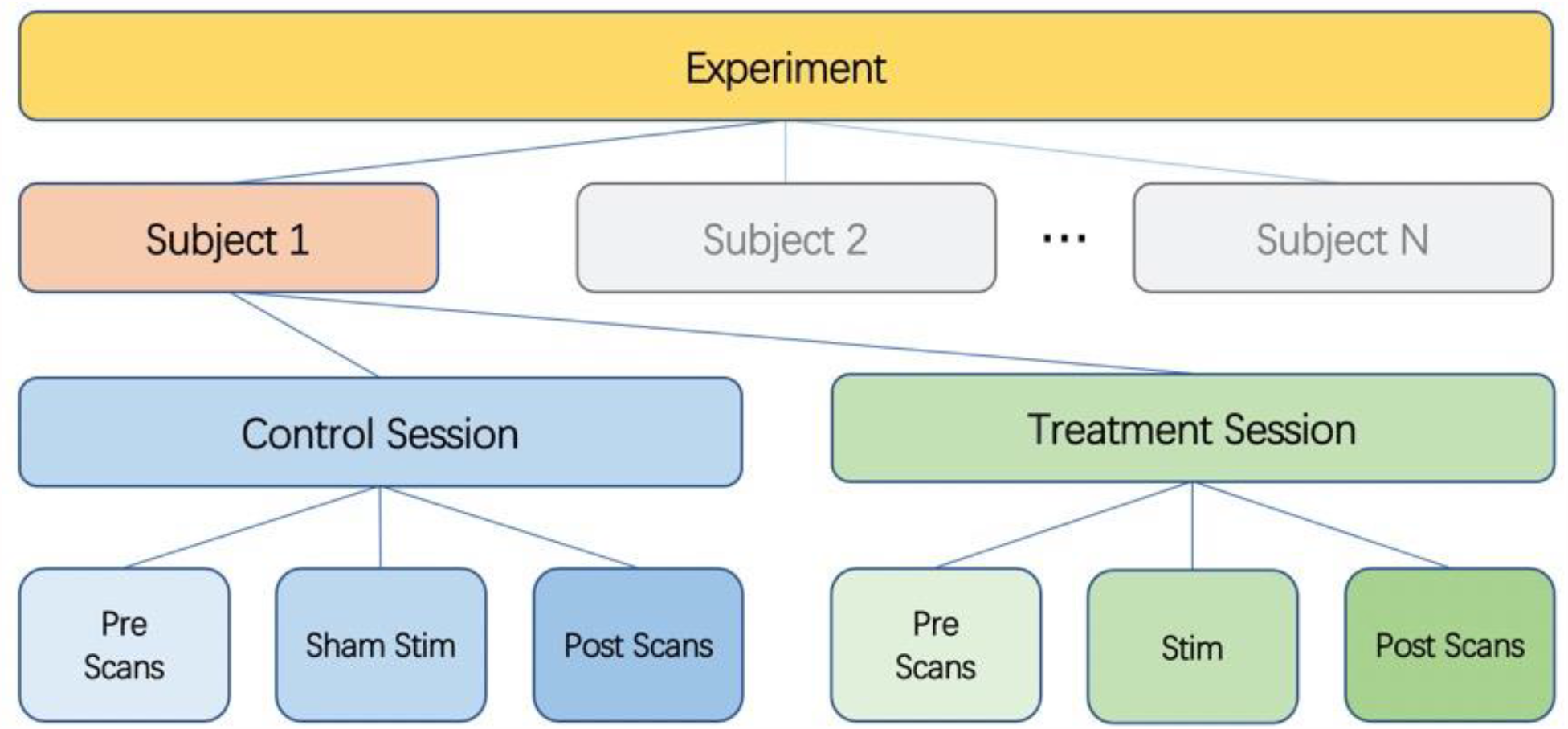
Diagram of experimental design. Each subject undergoes a control session and a treatment session. Each session consists of pre scans, a stimulation (sham for control session) section, and post scans.

All scanning visits consisted of a pre scan section, a stimulation (or sham) section, and a post scan section. For the first 6 subjects, the subjects wore the stimulation earpieces throughout the entire session, and the stimulation section took place while they remained lying inside the scanner. To improve the subject experience, the protocol was slightly modified for the subsequent 21 subjects. These 21 subjects wore regular earplugs during the MRI scans, came out of the scanner for the stimulation section, and then returned to the scanner with regular earplugs to finish the post scan section.

### MRI scans

Scans were acquired on a GE Discovery MR750 3.0T system. Pre and post scan sections consisted of (1) a high-resolution anatomical scan (MPRAGE, resolution=1mm^3^, 208 slices, FOV=25.6cm, TR=2500ms, TE=2.92ms, FA=8deg), (2) resting-state BOLD fMRI scans, and (3) arterial spin labeling (ASL) scans (2D pseudo-continuous ASL (PCASL), single-shot spiral, 100 repetitions, resolution=3.75mm×3.75mm, slice thickness=6mm, matrix size=64×64×24, FOV=24cm, TE=3.2ms, TR=4300ms, labeling duration=1800ms and post-labeling delay=1800ms) [26]. Only the anatomical and ASL scans are analyzed here. The resting-state BOLD scans will be analyzed in a separate paper. During all ASL scans, subjects rested with their eyes open and gently focused on a fixation point.

### taVNS

taVNS was delivered using custom-fit ear pieces designed by Nēsos®, (Redwood City, CA). Hydrogel electrodes were placed on the ear piece surface interfacing with the cymba concha. Alcohol wipes were used to clean the cymba concha prior to inserting the ear pieces. Stimulation was applied to the left ear, targeting vagal afferents in the cymba concha based on known vagus nerve anatomy [5,27,28]. For all visits, the stimulation (or sham) section took place between the pre and post scan sections.

On both scan days, a sensation test was performed to determine a subject’s threshold for perceiving the stimulation, which entailed stimulating at 1.0 mA and increasing by increments of 0.5 mA (up to a maximum of 5.0 mA). This procedure was repeated until the subject reported feeling the stimulation at the same level three times. The sensation test was followed by a 15-minute stimulation period. The stimulation was delivered continuously at 20 kHz, using biphasic square waves with pulse width of 20us. The stimulation amplitude was set at 75% of a subject’s perceptual threshold (1.5-3.8 mA; impedance 3-9.5 kΩ at 20 kHz) on the “active” stimulation day and was set to zero on the control day (“sham” condition). The order of “active” and “sham” stimulation days was randomized across subjects.

Since the stimulation was delivered at 75% of perceptual threshold, subjects were blinded to whether they were undergoing the sham or stimulus treatment. At the end of their second session, all subjects were asked whether they could identify their control and treatment sessions across their two visits.

### Data Processing and Analysis

Data from the ASL scan were volume registered and quantified into physiological CBF units (mL/100g/min) via local tissue correction with a proton-density scan acquired during the same scan section [29]. The SCORE algorithm [30] was used on a per-slice basis to censor out outlier timepoints in the CBF time series before computing the average CBF map. Using AFNI software [31], the high-resolution anatomical scans were transformed into MNI space, and the CBF volumes were then warped to MNI space using the transformation matrices from the anatomical transformation, with a final voxel resolution of 4mm^3^.

For group analysis in MNI space, Post-Pre CBF difference maps were computed for control and treatment sessions. A two-sided paired t-test was performed for the main contrast (Post-Pre)_treatment_ - (Post-Pre)_control_ on a per-voxel basis using AFNI 3dttest++. This contrast is equivalent to the interaction effect for a two-way (Post, Pre by Treatment, Control) repeated measures ANOVA. Clustering thresholds were estimated using permutation-based methods on the residual data via the -Clustsim option in 3dttest++. A voxelwise threshold of p<0.01 and a cluster size threshold of 22 voxels (clusters defined with voxel faces touching) were used to assess results at a FWE-corrected p<0.05 level [32].

Post-hoc two-sided paired t-tests were also performed for each session (control and treatment), and were assessed at a voxelwise threshold p<0.01 and a FWE-corrected p<0.05 level (using cluster size thresholds of 35 and 51 voxels for the control and treatment contrasts, respectively). In addition, post-hoc analyses were performed using the mean CBF values of any significant clusters found from the main contrast of (Post-Pre)_treatment_ - (Post-Pre)_control_.

## Results

### Participants and Stimulation

Out of twenty-seven total subjects, two dropped out after their first session, due in part to the COVID-19 related suspension of research activities during the course of the study. The remaining twenty-five subjects completed both the control and treatment sessions. Five subjects were excluded from the study due to technical errors or severe image artifacts in at least one of their CBF scans, leaving twenty total subjects included in the analysis. These twenty subjects received a mean stimulation amplitude of 3.24 ± 0.56 mA (min = 1.5mA, max = 3.8mA). The control and treatment sessions were matched to take place at the same time each day whenever possible, with an average time-of-day difference of 0.9 ± 1.32 hours (min = 0 hours, max = 4 hours).

When asked whether they could identify the control and treatment sessions across their two visits, sixteen of the twenty subjects responded that they could not guess. Of the remaining four subjects who indicated they could identify their sessions, three guessed correctly and one guessed incorrectly. Overall, an average of 24.81 ± 6.93 minutes elapsed from the end of the stimulation to the start of the post section ASL scan for these subjects.

### Group-level whole-brain analysis

For the main contrast shown in Figure 2a, application of clustering analysis at voxelwise p<0.01 and FWE-corrected p<0.05 revealed one significant cluster spanning both hemispheres of the posterior cerebellum (71 voxels, peak MNI coordinate [-6, 78, −36]). Figure 2b and 2c show session-level contrasts that support the main cluster result. Figure 2b shows no significant CBF changes in the control session, while Figure 2c shows a left hemispheric cerebellar cluster (69 voxels) and a right hemispheric cerebellar cluster (74 voxels) with significant CBF decreases in the treatment session.

**Figure 2:**
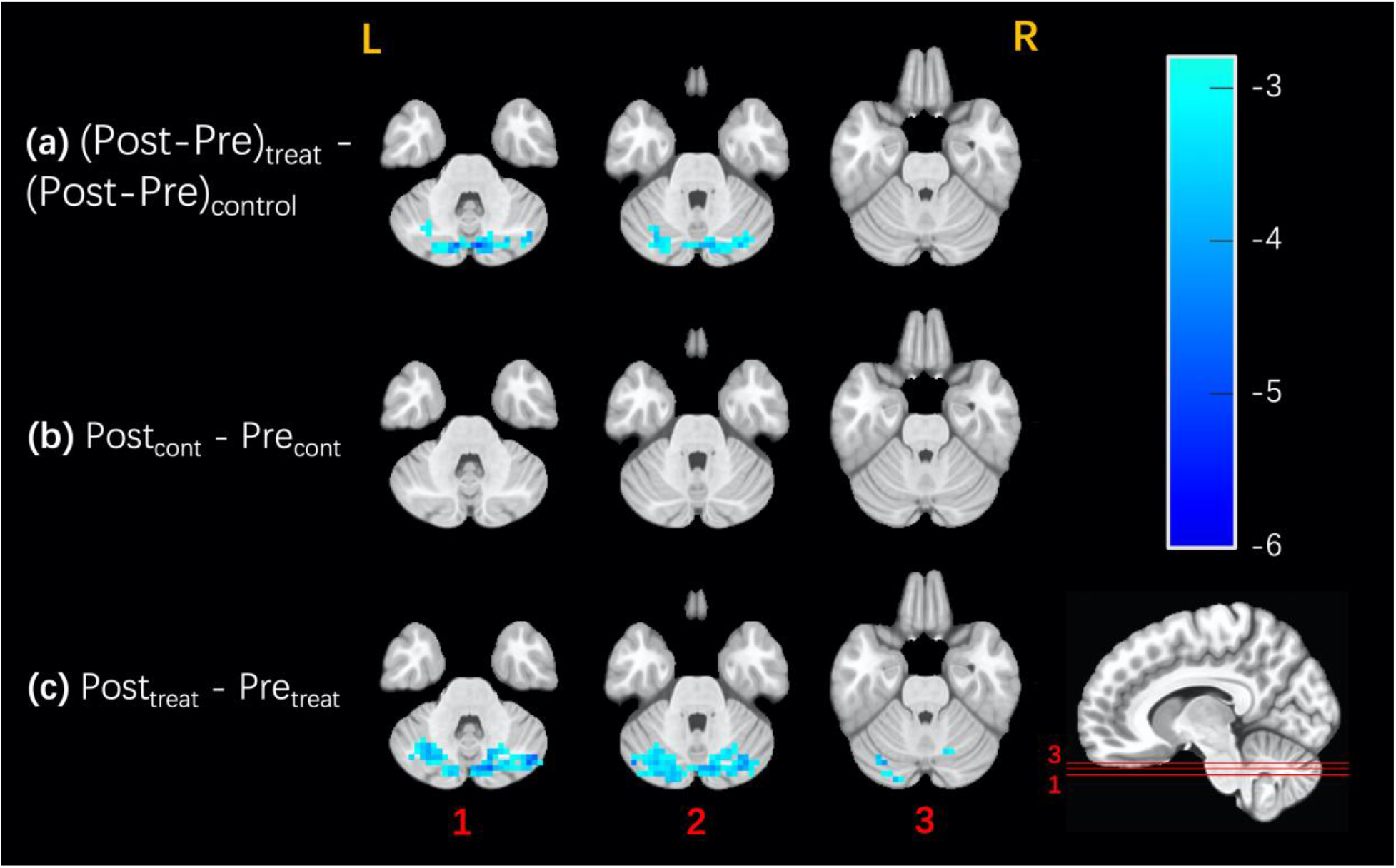
Maps showing clusters of voxels with significant effects, FWE-corrected at p<0.05 with voxel-wise threshold p < 0.01. Contrast-dependent cluster size thresholds based on permutation testing are indicated below. Surviving voxels are color-coded by their t-statistic value and overlaid on anatomical images, with the same slices shown for all three contrasts. No significant clusters were observed in any of the other slices. **(a)** For the main (Post-Pre)_treatment_ - (Post-Pre)_control_ contrast, one cluster with 71 voxels survived with a cluster size threshold of 22 voxels. **(b)** For the Post_control_ - Pre_control_ contrast, no clusters survived with a cluster size threshold of 35 voxels. **(c)** For the Post_treatment_ - Pre_treatment_ contrast, two clusters survived with a cluster size threshold of 51 voxels.

We used the probabilistic atlas of the cerebellar lobules as defined in [33,34] and the AFNI whereami function to determine the overlap of the significant clusters with the cerebellar lobules. For the main contrast, the single cluster showed 96.3% overlap with the cerebellar lobules, with 47.2% of the cluster overlapping with left and right Crus I, 44.8% with left and right Crus II, 3.5% with Vermis Crus II, and 0.8% with left and right lobules VIIb. For the treatment contrast, the left hemispheric cluster had 94% overlap with cerebellar lobules, with 75.5% overlap with left Crus I, 15.8% with left Crus II, and 2.6% with left lobule VI. The right hemispheric cluster had 89.7% overall overlap consisting of 45.4% overlap with right Crus I, 28.5% with right Crus II, 8% with right lobule VI, 3.3% with right dentate, 2.7% with Vermis Crus II, 1.8% with left Crus II, and 0.1% with right VIIb.

Post-hoc analyses of the mean CBF values from the main contrast cluster are shown in Figure 3. The cluster shows a significant decrease in CBF related to treatment (p<0.001, −15.90%) alongside a statistically insignificant increase in the control session (p=0.19, +4.02%). Consistent with the maps shown in Figure 2, the post-hoc analyses indicate that the contrast of interest mainly reflects significant treatment-related decreases in cerebellar CBF.

**Figure 3:**
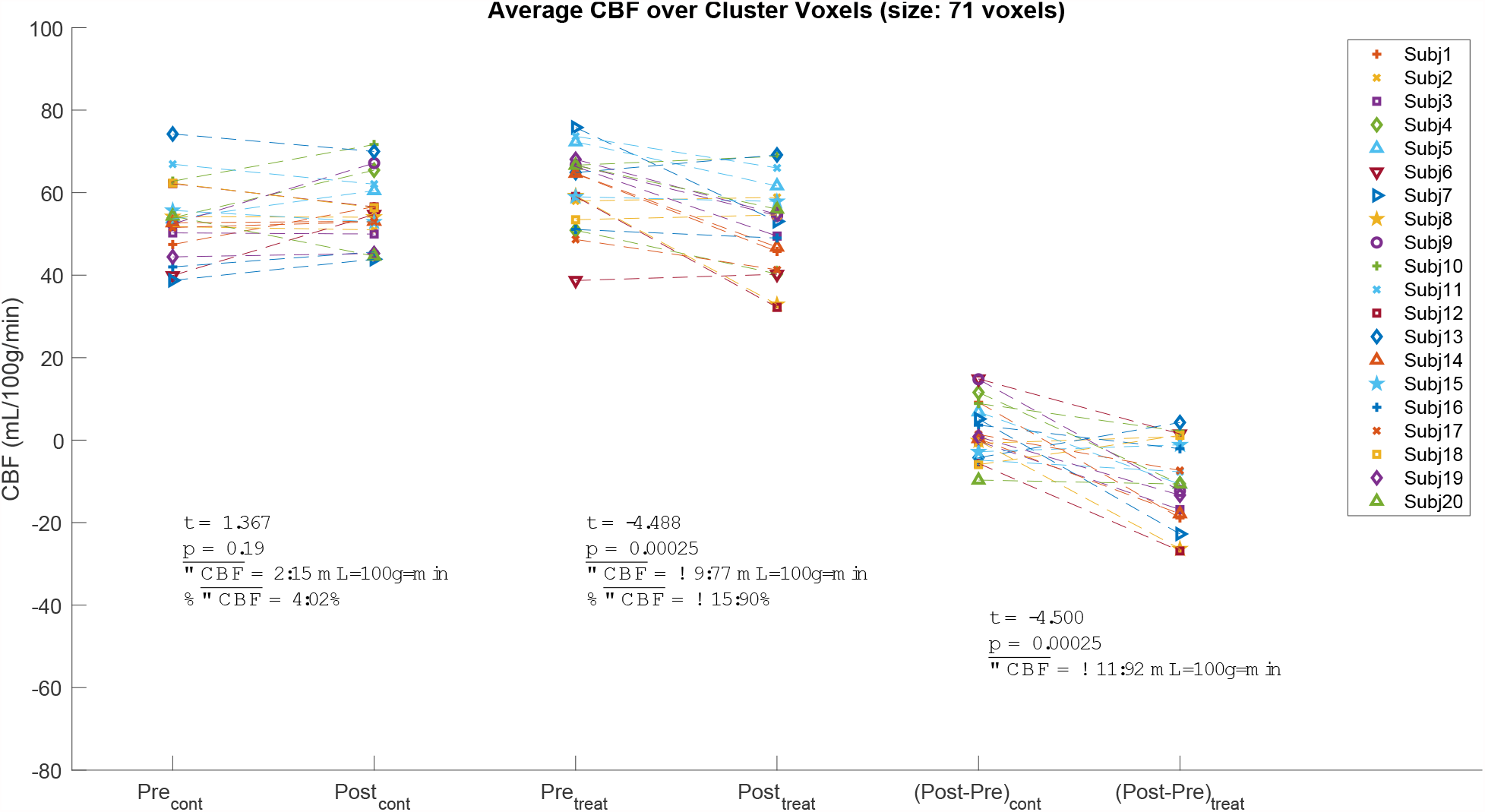
Post-hoc analysis with mean per-subject CBF values from the cluster shown in Figure 2a. The Pre and Post values from the control and treatment sessions are shown in the leftmost and center pairs, respectively. The (Post-Pre) CBF differences from the control and treatments sessions are shown in the rightmost pair. There was a significant treatment-related decrease in CBF (p < 0.001, −15.90%) alongside a weak, statistically insignificant increase in the control session (p = 0.19, +4.02%), resulting in a significant main effect (p<0.001).

## Discussion

Our findings reveal a significant bilateral decrease in cerebellar CBF as a result of 15 minutes of sub-threshold taVNS, reflecting physiological changes that persisted for many minutes after the treatment. These effects were observed using a stimulus paradigm and duration similar to those of the daily dose employed in the recent open label clinical trial of [8].

In contrast to the current findings, prior studies have reported both unilateral [24] and bilateral [19,21,22] increases in cerebellar CBF. However, there are several key methodological differences that are worth noting. First and foremost, the prior studies examined evoked CBF changes in response to the concurrent application of 30 to 90 seconds of VNS, whereas the current study assessed resting-state CBF measures obtained before and after the application of taVNS. Thus, the prior findings are limited to characterizing the acute cerebellar CBF response to VNS, while the current findings offer evidence of a sustained decrease in cerebellar CBF in reponse to taVNS. Second, the use of a sub-threshold high frequency (20 kHz) stimulus in the current study enabled the use of a single-blind placebo controlled design, in contrast to the open-label design with the super-threshold low frequency (20 to 30 Hz) stimuli employed in prior studies. The placebo control in the current study enabled a clean isolation of the direct vagal stimulation effects from those associated with stimulus-associated sensations and responses, a separation that was not possible with prior designs. Finally, the current findings were obtained in healthy subjects who were naïve to taVNS, whereas the prior results characterized the responses in MD or epilepsy patients at a time point that was hours to months after the commencement of continuous VNS therapy. Thus, the current study provides critical information about the effect of taVNS on the healthy brain that can serve as a baseline for the future assessment of effects in clinical populations.

Stimulus-induced changes in cerebellar activity have also been observed with BOLD fMRI, but the experimental methods and results greatly vary across studies with reports of both evoked activations in healthy subjects [12–14] and epileptic patients [35] and deactivations in healthy subjects [36] and MD subjects [37,38]. As the BOLD signal exhibits a complex dependence on neural, vascular, and metabolic factors, a definitive link between observed BOLD changes and underlying CBF changes cannot be established without additional experimental measures [39]. However for most conditions, task-evoked increases and decreases in the BOLD signal are generally thought to reflect concomitant CBF increases and decreases. Thus, while the fMRI findings cannot resolve whether VNS induces increases or decreases in cerebellar CBF, they offer general support for the potential of VNS to modulate CBF levels in the cerebellum.

In this study, significant CBF changes were limited to the cerebellum. The prior PET and SPECT studies reported CBF changes in additional brain regions, although the results were highly variable across studies. As noted above, a number of these studies considered the evoked CBF response as compared to the sustained response reported here. Two SPECT studies examined the effects of 4 to 10 weeks of chronic VNS [17,18] and reported increases and decreases in CBF in multiple brain regions, but there were differences between the two studies in the regions that were identified and neither study reported cerebellar CBF changes. In light of the variability observed in prior studies and the significant methodological differences discussed above, discrepancies between the prior and current findings are to be expected. In particular, the interpretation of the changes observed in the prior studies typically discussed the potential modulation of activity in regions that were thought to be associated with the disease state. These brain regions would not necessarily show an effect in healthy subjects. Nevertheless, a future study examining the effects of sub-threshold taVNS over a longer period (e.g. weeks) would be of interest to determine if CBF changes can be detected in areas beyond the cerebellum.

While the cerebellum has traditionally been viewed as a motor control region, a wealth of evidence suggests that it also plays important roles in cognitive, affective, and social processing [40–42]. In particular, the cerebellar regions Crus I and Crus II, which exhibited the greatest overlap with significant CBF changes in this study, are considered to be primarily associated with non-motor functions, such as working memory, executive function, language processing, social mentalizing, and emotional cognition [41,43,44]. Interestingly, the cerebellum has a purported role in conditions for which taVNS has beneficial effects. For example, cerebellar dysfunction has been implicated in major depressive disorder (MDD) [45], and a preliminary clinical study found that a multi-week regime of taVNS (relative to sham) can alleviate depression severity as assessed via the Hamilton Depression Scale in MDD patients [1]. TaVNS-induced cerebellar activation has also been reported in an fMRI study of MDD patients, though individual differences in activation were not related to clinical outcomes [46]. Furthermore, taVNS has shown promise as a treatment for migraine [10], another disorder that has been linked to functional and structural alterations within the posterior cerebellum, namely Crus I and Crus II [47,48]. Finally, a recent study found that a 12-week regime of taVNS treatment (with parameters similar to those used in this study) improved symptoms for patients with rheumatoid arthritis (RA) [8]. Reduced gray matter of the cerebellum has been associated with higher levels of peripheral inflammation in RA patients [49]. Together, these findings suggest that the link between taVNS therapeutic efficacy and cerebellar function should be further explored.

In conclusion, the present results indicate that 15 minutes of high-frequency sub-threshold taVNS can induce sustained CBF decreases in the bilateral posterior cerebellum in a sample of healthy subjects. Further work is needed to determine if similar changes can be observed in clinical populations and to assess if the measures can provide greater insight into the mechanisms underlying treatment outcomes. In addition, future research should also address how stimulation parameters such as frequency, duration, and electrode location impact the response.

## Conflicts of Interest

Funding for this study was provided by an unrestricted grant from Nēsos^®^ (Redwood City, CA), which also provided hardware support for this study. Kelly H. MacNiven and Gary Heit report consultant fees from Nēsos. Konstantinos Alataris, Gary Heit and Vivek Sharma report equity holdings in Nēsos. Konstantinos Alataris and Vivek Sharma are current employees. Thomas Liu reports grants from Nēsos. All other authors declare no competing interests.

## Acknowledgements

This work was supported by a research grant from Nēsos Corporation, Redwood City, CA.

